# Drug-induced phospholipidosis as an artifact in antiviral drug repurposing

**DOI:** 10.1101/2025.11.20.689520

**Authors:** Isabella S. Glenn, Lu Paris, Alex D. White, Virginia G. Garda, Mauricio Montano, Mir M. Khalid, Aimee W. Kao, Melanie Ott, Brian K. Shoichet

**Affiliations:** Department of Pharmaceutical Chemistry, University of California San Francisco (UCSF), San Francisco, California 94143, United States; Weill Institute for Neurosciences, Memory and Aging Center, Department of Neurology, University of California San Francisco (UCSF), San Francisco, California 94158, United States; Small Molecule Discovery Center, Department of Pharmaceutical Chemistry, University of California San Francisco (UCSF), San Francisco, California 94143, United States; Gladstone Institutes, San Francisco, California 94158, United States; Department of Medicine, University of California, San Francisco, San Francisco, California 94158, United States; Chan Zuckerberg Biohub, San Francisco, California 94158, United States

## Abstract

Drug repurposing in principle can speed antiviral drug discovery. Among the molecules most frequently advanced in such repurposing efforts are a group of structurally diverse cationic amphiphilic drugs (CADs). While CADs have shown micromolar to mid-nanomolar antiviral activity in cell-based assays, they can induce phospholipidosis, confounding in COVID-19 repurposing. A barrier to the identification of phospholipidosis inducers has been the involved nature of the microscopy assays used to characterize them. To ease the identification of these artifacts, we describe a rapid microplate-based assay to detect phospholipidosis. Leveraging this assay, we quantified the prevalence of phospholipidosis-inducers across several cell-based antiviral repurposing screens. We selected 40 drugs reported to have micromolar antiviral activities and found that 26 of them (65%) induced phospholipidosis within the same concentration range as their reported antiviral activities. Intriguingly, we identified four non-CADs that also induce phospholipidosis, revealing a new group of drugs that can lead to this toxic event and highlighting the importance of facile experimental assays to detect it. Understanding how phospholipidosis can confound antiviral drug discovery, and its rapid detection, will help prevent what is an apparently general artifact, active across viruses, from distracting investigators from potentially more useful candidates.

## Introduction

Drug repurposing is a popular strategy to identify new indications for approved or investigational drugs^1^. Developed drugs are de-risked molecules, and advancing them eliminates many of the costly and time-consuming pre-clinical steps in drug discovery and development^2^. With the Covid-19 pandemic, drug repurposing took on greater urgency, given the need to rapidly discovery new antiviral therapeutics^3^. Indeed, two of the triumphs of Covid-19 drug discovery, nirmatrelvir and remdesivir, both reflect some aspect of drug repurposing—the first drawing on extensive efforts in developing Mpro inhibitors for SARS-1^4, 5^, the second emerging from a drug discovery effort against ebola virus^6, 7^. Still, these drugs and their antecedents were themselves antivirals with defined mechanism of action. During the pandemic, many investigators cast a wider net, testing large drug libraries, most of which targeted human proteins, seeking molecules that would be fortuitously^8–10^ or by mechanism ^11–14^ active against the virus. While several promising molecules emerged from this effort, many of the repurposed molecules, despite having low µM to even mid-nM activities in cell culture, were ultimately shown to be false-positives acting via artifactual mechanisms^15, 16^. Among the most prominent of these is drug-induced phospholipidosis (DIPL)^17^.

Phospholipidosis is a lysosomal storage disorder characterized by the intracellular accumulation of phospholipids, as occurs in some pathologies such as Niemann-Pick disease^18^. A morphological-hallmark of phospholipidosis is the formation of lamellar bodies and “foamy” and “whorled” membranes that can be captured by electron microscopy ^19^. The effect is of wide concern in pharmaceutical development, where drug-*induced* phospholipidosis (DIPL) can cause idiopathic toxicities, often emerging late in development, in sensitive organs including the liver and kidney^20, 21^. This has motivated the development of fluorescently conjugated phospholipids, such as NBD-PE, to monitor phospholipid accumulation by microscopy^22, 23^.

Chemically, phospholipidosis inducers are diverse, and belong to drug classes including antibiotics, antidepressants, antipsychotics, and antiarrhythmics ^18^. Physically, phospholipidosis inducer share several properties: most are cationic at physiological pH and are relatively hydrophobic, with cLogP (calculated log of the octanol:water partition ratio) values often > 3 (and so are called “cationic amphiphilic drugs” (CADs))^24^. CADs become further ionized in the acidic lysosomal environment, where they accumulate. This lysosomal accumulation is thought to be a driving force for DIPL, though a full mechanism has yet to be elucidated^21^.

Many of the drugs repurposed for activity against SARS-CoV-2 in cell culture were CADs, and their antiviral activity was subsequently found to correlate with the phospholipidosis they induced^17^. In the same study, drugs that were cationic but not amphiphilic did not induce phospholipidosis and were not antiviral, while known CADs but were unknown to be antiviral could be shown to have anti-proliferative effects against SARS-CoV-2 in cell culture. Whereas the CADs could have anti-viral effects down to the 100 nM range, they could not be optimized beyond that, consistent with their phospholipidosis range of activities. Exactly how drug-induced phospholipidosis translated into apparent antiviral activity in cell culture remains to be fully characterized, though it is thought that the disruption of phospholipid homeostasis affects the ability of SARS-CoV-2 to replicate, perhaps via disruption of the double-membrane vesicles on which it depends. Meanwhile, the antiviral activity of the CAD phospholipidosis-inducers did not translate into animal efficacy but only into the toxicity with which phospholipidosis inducers are associated with at high concentrations^17, 25^ ^26^.

Here we investigate two further impacts of phospholipidosis on antiviral repurposing: its role on drugs repurposed against non-SARS-CoV-2 viruses, and its role in drugs outside of the traditional cationic amphiphiles with which it has been most associated. To do so, we adopted a new high-throughput assay to quantify DIPL. We used this assay to investigate 40 drugs that had been reported as candidates for antiviral repurposing across seven viruses^27–45^, including several that were not cationic and so were not expected to induce phospholipidosis by the usual metrics^46, 47^.

Implications for drug repurposing, the range of molecules that might induce phospholipidosis in drug development, and the ability to rapidly test for this effect will be considered.

## Results

### A rapid assay to detect drug-induced phospholipidosis

A rapid method to detect phospholipidosis is crucial to ruling its confounding effects. Electron and confocal microscopy are the gold-standard for measuring phospholipid accumulation but are slow and require access to imaging facilities. In the initial study of drug-induced phospholipidosis (DIPL) among SARS-CoV-2 inhibitors, a microscopy-based NBD-PE assay was used that required solubilizing NBD-PE in ethanol, followed by extended sonication to ensure proper dispersion in solution. Cells were typically fixed before imaging and although the experiments were conducted by high-content microscopy, processing the imaging datasets was time consuming and required both specialized software and expertise. Though widely used in pharmaceutical research to identify DIPL, this NBD-PE assay, too, is unsuited for rapid detection. Accordingly, we adapted an assay first introduced by a group at NCATS^48^. While this assay still utilizes high-content microscopy, LipidToxRed is used in place of the traditional NBD-PE reagent. LipidTox Red offers several advantages: it comes as a ready-mix solution bypassing solubility challenges of traditional PLD reagent, is suitable for live-cell imaging, and is more sensitive for PLD detection compared to NBD-PE ^49^. Further, we describe a modified version of this assay that can be performed using a microplate reader, reducing image processing time and improving accessibility to non-experts (**Figure 1A**).

**Figure 1.**
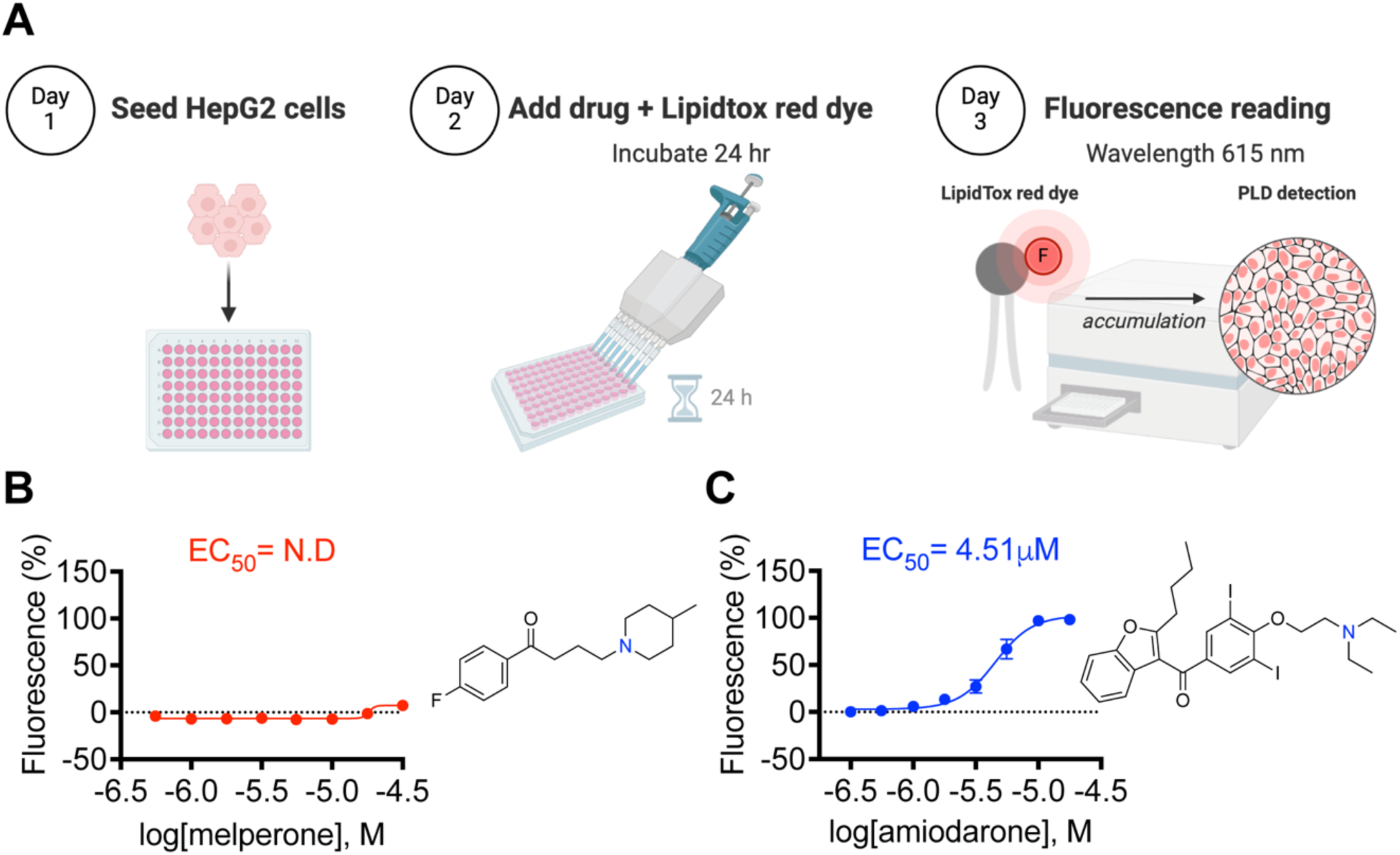
Overview of the phospholipidosis assay. (**A**) Schematic of the three-day experiment. (**B**) Results for the non-PLD inducing cationic drug melperone and (**C**) the well-characterized CAD PLD inducer amiodarone. All data shown was measured in Hep G2 cells using LipidTox Red. Error bars represent SD for three independent experiments performed in triplicate.

Consistent with earlier studies^48^, the plate-based assay performed well against positive and negative controls, accurately distinguishing a non-PLD inducing cationic drug melperone (**Figure 1B**) from the well-characterized PLD inducer amiodarone (**Figure 1C**). Encouragingly, the potency of amiodarone was consistent in the plate-based assay to previously reported values from orthogonal assays^17, 48, 50^, including our earlier microscopy-based study^17^. This assay can rapidly detect and quantify drug-induced phospholipidosis without lengthy microscopy experiments and analyses.

### Screening for phospholipidosis among drugs repurposed as antivirals

To identify antiviral repurposing candidates to screen for phospholipidosis, we searched the literature for drugs reported as antivirals in cell-based work with an unclear antiviral mechanism of action. We filtered these for those with similar physiochemical properties to known phospholipidosis inducers: clogP≥2 and calculated pKa ≥7.4. Forty compounds met these criteria (**Table 1, Table S1**), including four that did not pass both property cutoffs but that had multiple reports of antiviral activity. These were tested for induction of phospholipidosis using the assay introduced above. Of the 40 drugs, 26 (65%) induced PLD in the same potency range as their reported antiviral activity (**Figure 2A, Figure S2**). Of these, 22 were stereotypical CADs with top ten most potent PLD inducers ranging from 400nM to 5µM (**Figure 2B**).

**Figure 2.**
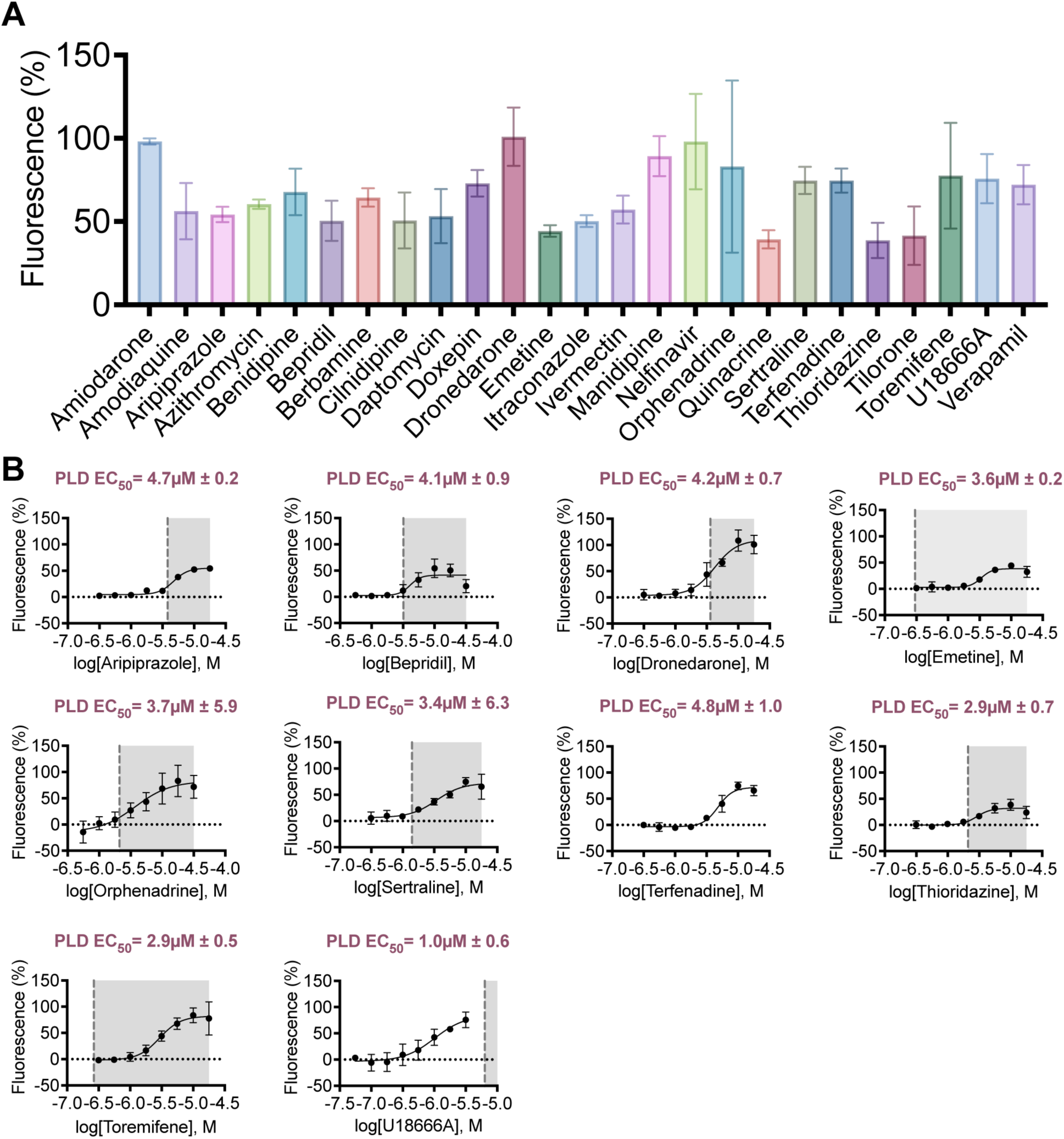
Repurposed drugs that induce phospholipidosis. (**A**) Percentage fluorescence with treatment of drug at a single-point concentration at approximately PLD E_max_. Itraconazole and U18666A were screened at 3.16µM and Daptomycin, Nelfinavir, Tilorone, and Verapamil at 31.6µM. All other repurposed drugs were screened at 17.8 µM. (**B**) Ten most potent phospholipidosis inducers that are prototypical CADs with their corresponding literature reported antiviral EC_50_ to E_max_ boundaries colored in gray. All PLD data shown was performed in HepG2 cells using LipidTox Red phospholipid reagent with error bars representing SD for three independent experiments performed in triplicate. Data was normalized to positive control compound, amiodarone, and cell count using Hoechst nuclei stain.

**Table 1.**
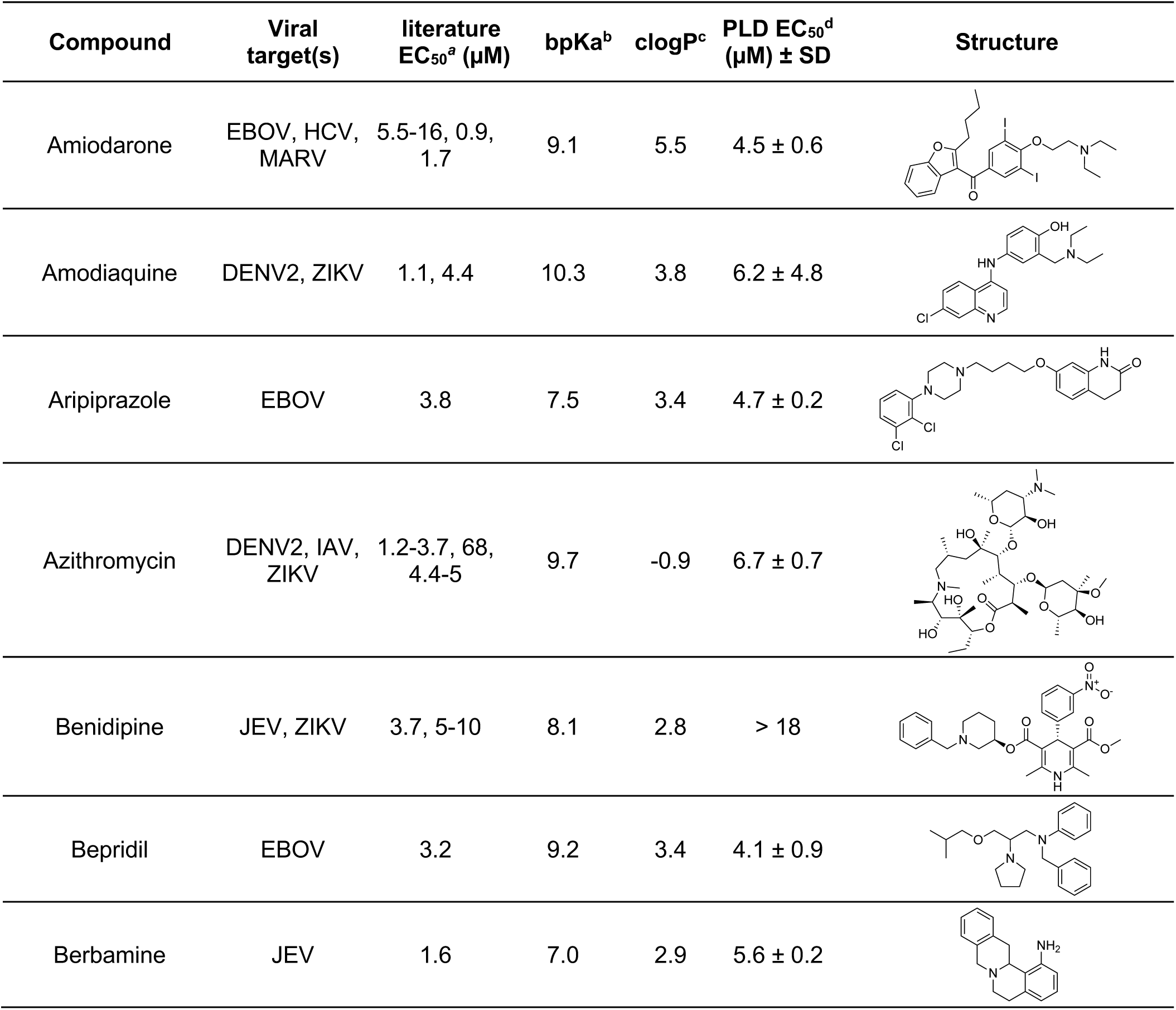
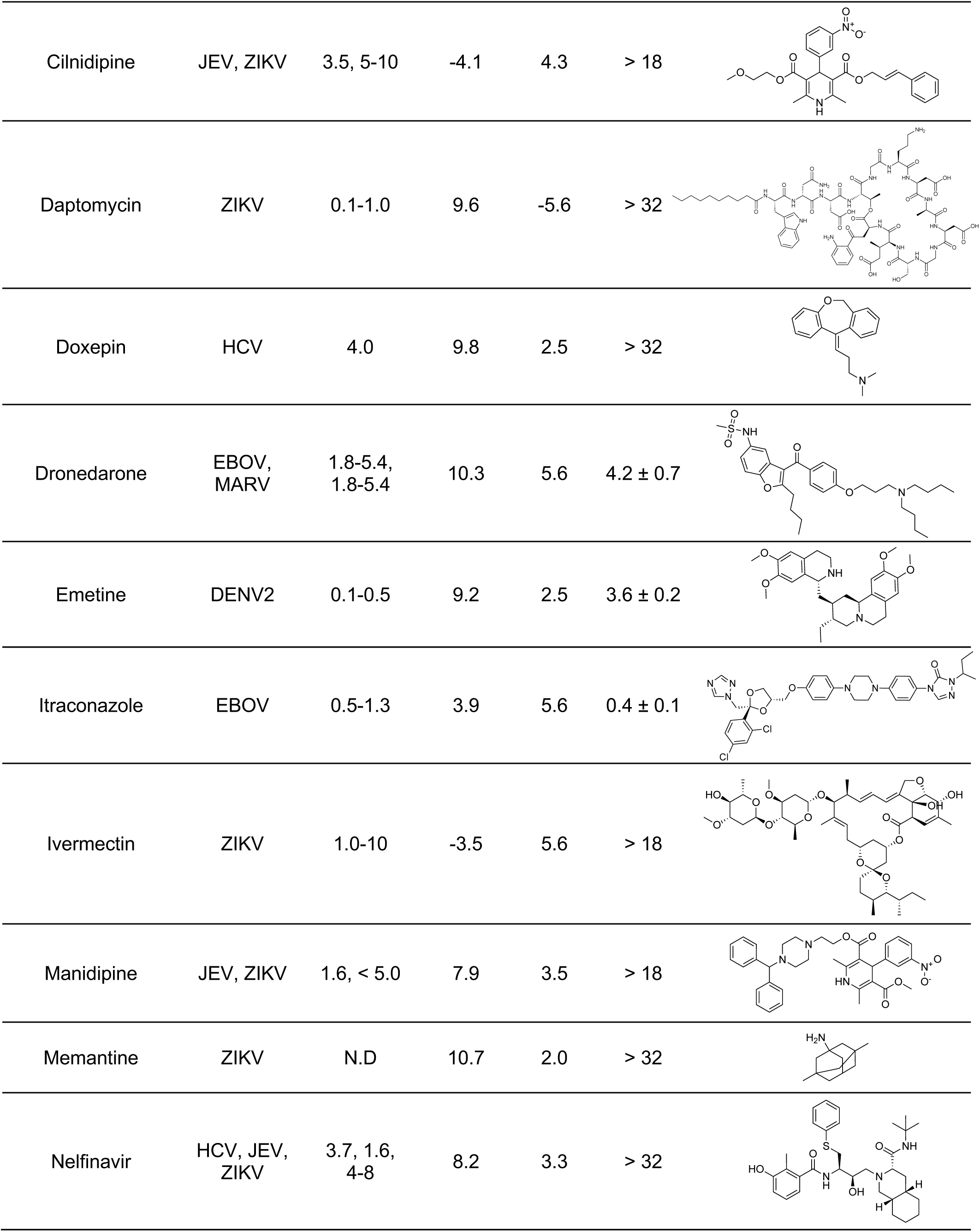
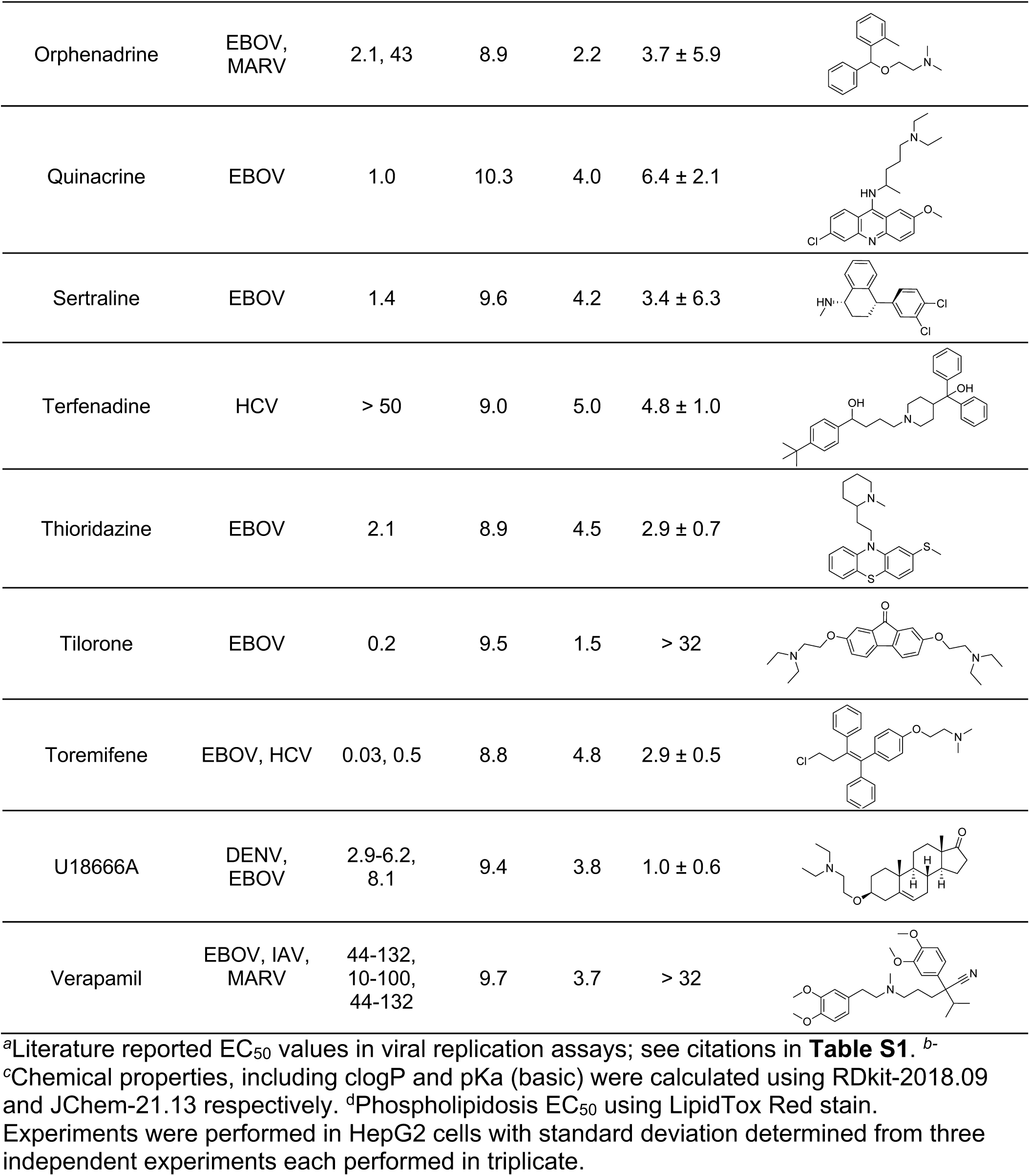
Repurposed drugs that induce phospholipidosis.

Intriguingly, four compounds that did not meet either the clogP or pKa cutoffs and would not ordinarily be classified as CADs did induce phospholipidosis (**Figure 3**). These included two neutrally charged PLD inducers, itraconazole and ivermectin (**Figure 3C**), and two highly soluble cationic drugs that had cLogP values of −0.9 and −5.6: azithromycin and daptomycin, respectively (**Figure 3B**). The four compounds were initially included in the screen based on their frequent appearance in the antiviral literature and lack of mechanism. Thus, while status as a CAD is the best predictor of being a phospholipidosis-inducing drug, it is neither necessary nor sufficient to predict such activity, a point also made by other studies^51^ and to which we will return.

**Figure 3.**
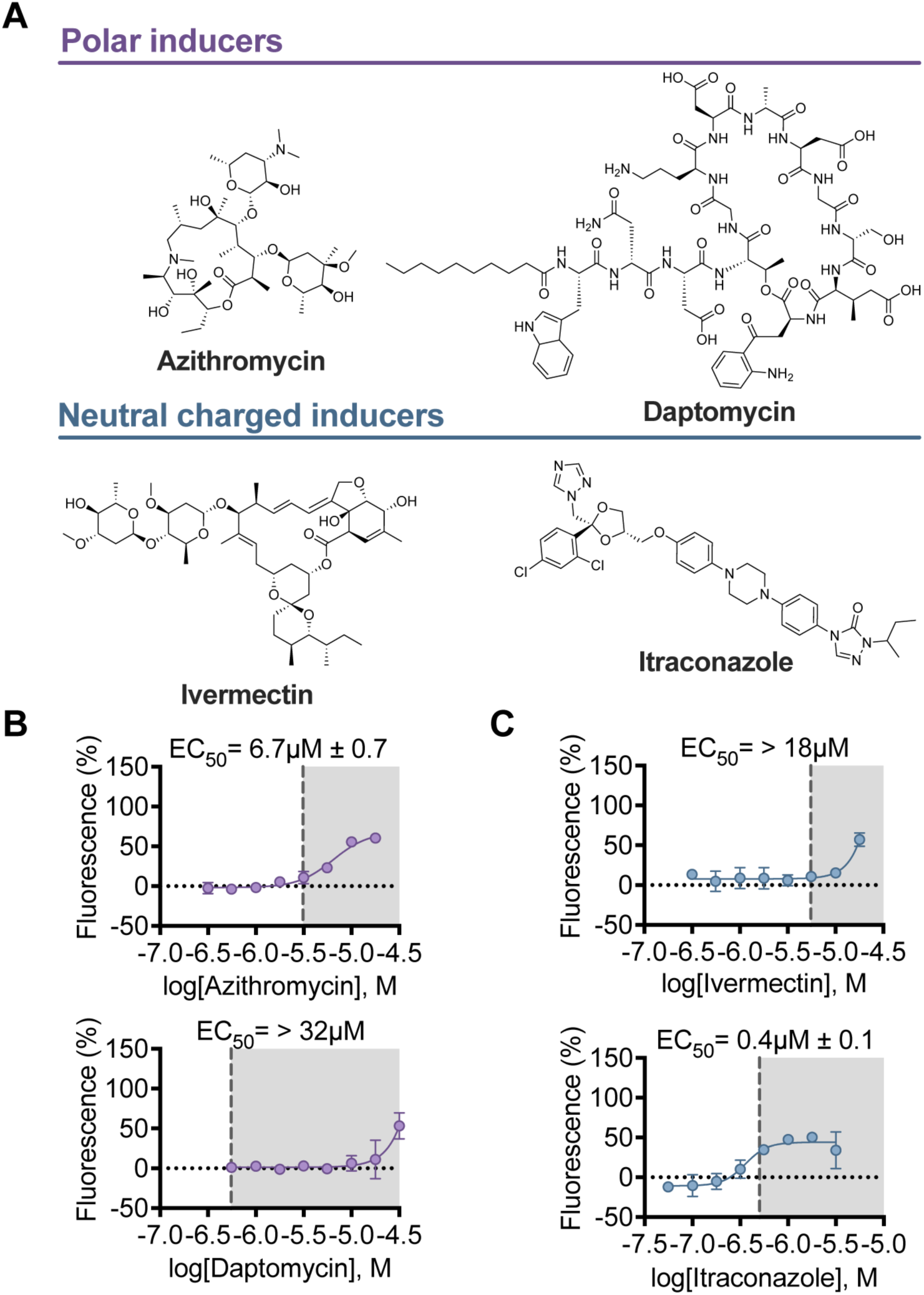
Non-CAD phospholipidosis inducers. (**A**) Chemical structures of non-CAD inducers. (**B**) More polar phospholipidosis inducers clogP< 2 and (**C**) Uncharged phospholipidosis inducers. All data shown was performed in HepG2 cells using LipidTox Red phospholipid stain. Error bars represent SD for three independent experiments performed in triplicate. Data was normalized to positive control compound, amiodarone, and cell count using Hoechst nuclei stain.

To ensure that our non-CAD compounds were indeed inducing phospholipidosis and not merely confounding the assay, we cross-validated some of our prior hits (**Figure 1**, **Figure 2**) and the non-CAD phospholipidosis inducers (**Figure 3**) using traditional NBD-PE dye and microscopy (**Figure 4**). We first validated that negative control compound melperone and the positive control amiodarone agreed with our results and literature reports (**Figure 4A**, **Figure 4B**). Next, we screened terfenadine (a prototypical CAD), itraconazole (non-cationic), and azithromycin (non-lipophilic) for phospholipidosis and found all three were inducers by the traditional assay (**Figure 4B**). The efficacy and potencies achieved after quantifying NBD-PE intensities (**Figure 4C**, **Figure 4D**) aligned with the data from our LipidTox red assay demonstrating the assay retains accuracy despite being higher-throughput.

**Figure 4.**
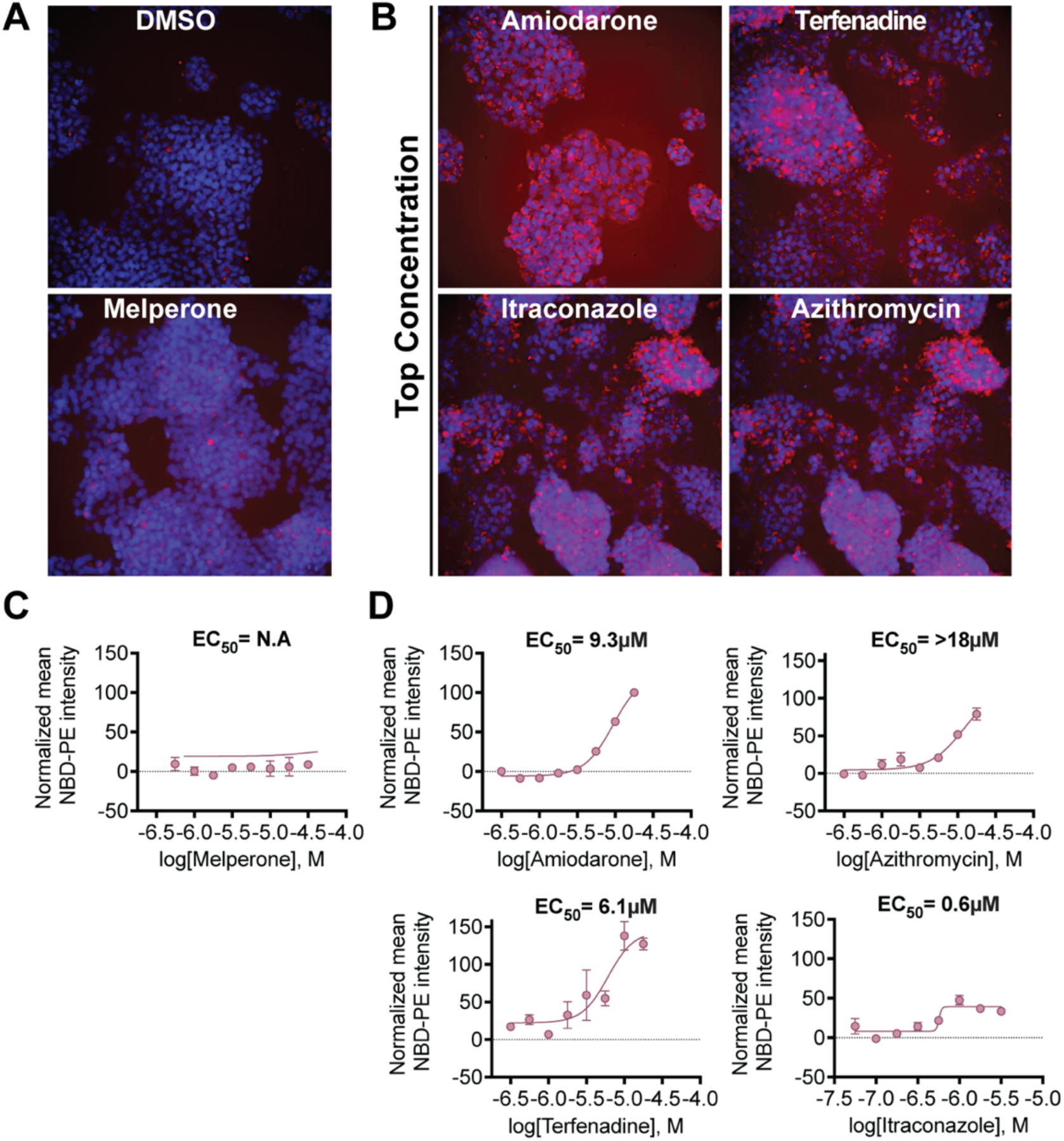
NBD-PE phospholipidosis assay. (**A**) Non-PLD inducing CAD melperone (31.6µM) and DMSO control (**B**) **Top panel** traditional CADs amiodarone (17.8 µM) and terfenadine (17.8 µM). **Bottom panel** non-CAD Itraconazole (pKa ≤ 7.4) (1.78 µM) and azithromycin (clogP< 2) (17.8 µM). (**C**) and (**D**) quantification of mean fluorescence intensity as a concentration-response. All data shown was performed in HepG2 cells using NBD-PE phospholipid stain in triplicate with error bars representing SD. Images were taken from 9-fields with representative image shown for each condition with scale bar (20 µm). Mean NBD-PE intensity values were normalized to positive control compound, amiodarone, and cell count using Hoechst nuclei stain.

### Investigating the role of lysosomal phospholipases in drug-induced phospholipidosis

To examine potential mechanisms underlying drug-induced phospholipidosis and its in vitro antiviral effect, we assessed if inducers could alter lysosomal pH or directly inhibit lysosomal phospholipases. One proposed mechanism of drug-induced phospholipidosis involves CADs becoming protonated at an ionizable amine in the lysosome’s acidic environment, leading to ion-trapping. In this view, once trapped in the lysosome, these “lysosomotropic” drugs accumulate and potentially raise lysosomal pH, inactivating acidic hydrolases including phospholipases^52–55^. Such lysosomal alkalinization has been implicated in antiviral activity, with several studies reporting that the lysosomal V-ATPase inhibitor, bafilomycin A1, acts as a potent antiviral through this mechanism^56–58^. Given these precedents and that some of the PLD inducers characterized here do not contain an amine ionizable at accessible pH values, we investigated whether these PLD inducers elevate lysosomal pH and thereby disrupt lysosomal homeostasis.

To do so, we used the FIRE-pHLy (Fluorescence Indicator REporting pH in Lysosomes)^59, 60^ biosensor to screen our non-CAD PLD inducers (azithromycin, daptomycin, itraconazole, and ivermectin), a panel of PLD inducers ranging in potency, a non-PLD inducing cationic drug (melperone), and bafilomycin A1 as a positive control (**Figure 5A**). At concentrations where they led to strong phospholipidosis, many of the PLD inducers had neither substantial nor statistically significant effects on the Fire-pHLy ratio fold-change, relative to the positive control bafilomycin, indicating that the pH of the lysosomes remained relatively unchanged after the 24 hour drug treatment. For instance, while bafilomycin changed the FIRE-pHLy ratio by 2.46-fold versus DMSO, potent PLD inducers like amiodarone and U18666A had little detectable and no significant effect on the Fire-pHLy ratio. There were other PLD inducers, like azithromycin, emetine, and terfenadine, that did show statically significant fold-changes versus baseline (**Table S2**), but these values only ranged from 0.33 to 0.57 and were far below the change observed for positive control bafilomycin A (**Figure 5A**). These results suggests that while some PLD inducers may raise lysosomal pH, the extent of alkalinization is modest and in most cases the lysosomal pH remains unchanged.

**Figure 5.**
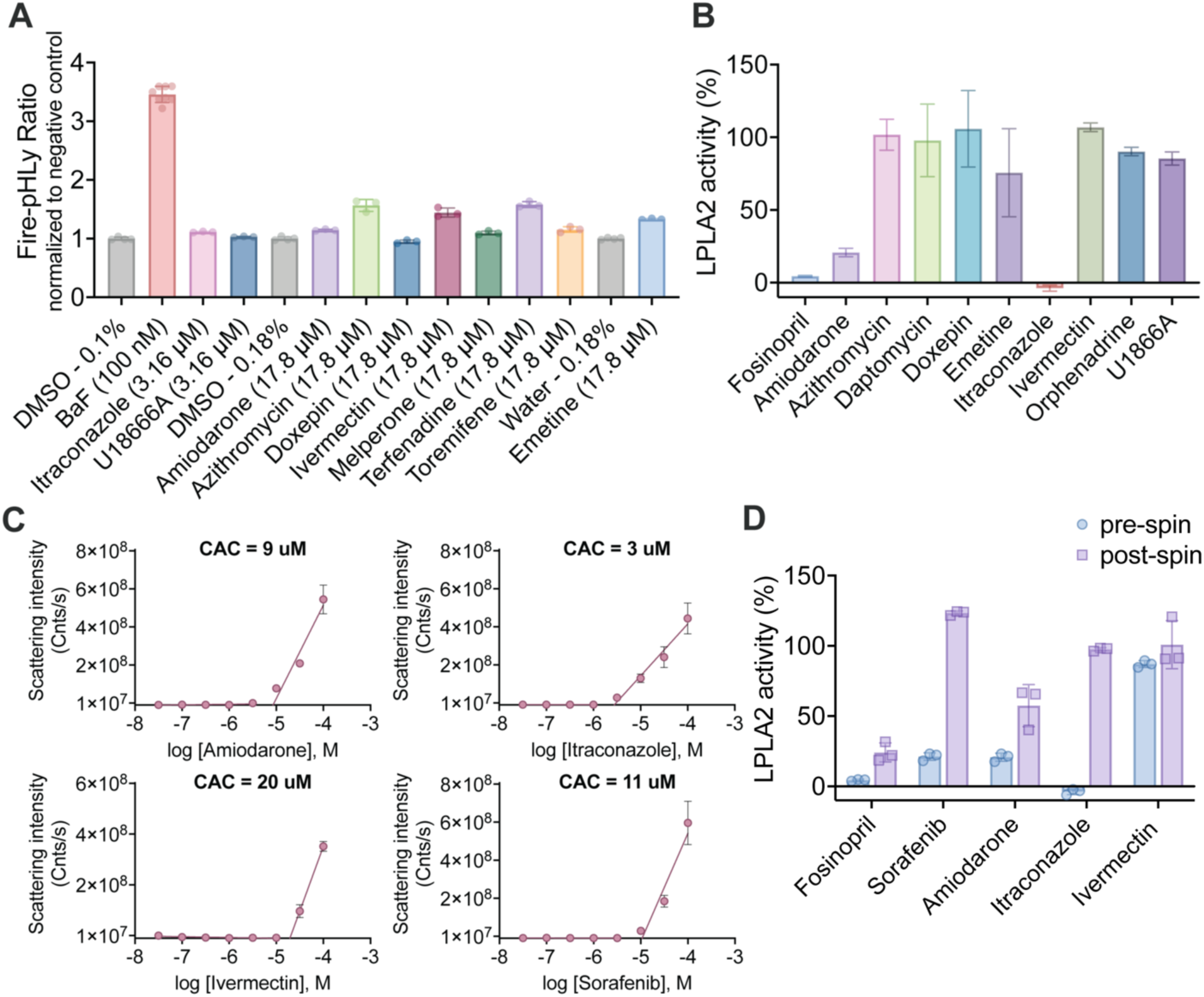
Phospholipidosis inducers effect on lysosomal pH and LPLA2 activity. (**A**) Bar graph quantification of FIREpHLy ratio fold-change in HepG2 cells after 24 hr drug treatment. Baseline control matched for vehicle concentration (DMSO or water) are shown to the left of each compound group. (**B**) LPLA2 activity assay. All compounds screened at 31.6µM. Data points are presented as mean ± SD from three technical replicates. (**C**) Dynamic light scattering performed on colloidal aggregator candidates by a scattering intensity threshold of 1 × 10^7^. The point of intersection serves as the critical aggregation concentration (CAC) value for each compound. All measurements were performed as three technical replicates in 50 mM KPi buffer. (**D**) LPLA2 activity before (blue) and after (purple) centrifugation screened at 31.6µM. Data points are presented as mean ± SD from three technical replicates

Next, we investigated whether PLD inducers could directly bind to and inhibit lysosomal phospholipases, as represented by lysosomal phospholipase A2 (LPLA2), a previously described target of PLD inducers^21, 61^. We tested several canonical CAD PLD inducers, our non-CAD PLD inducers, and characterized LPLA2 inhibitor, fosinopril, as a positive control (**Figure 5B**; we note that fosinopril is an anionic phosphonate and itself far from a CAD). While fosinopril potently inhibited LPLA2, as expected, amiodarone and itraconazole substantially inhibited LPLA2, but this was only observed at 31.6 µM, well-above the concentration that they induce PLD.

The relatively high concentrations necessary to inhibit LPLA2, and the high lipophilicity of amiodarone and itraconazole, prompted us to control for non-specific inhibition via colloidal aggregation, a common artifact among enzyme inhibitors^15^ and something to which we knew itraconazole was prone^62^. Colloidal drug aggregates act by sequestering enzymes^63^on their surface^64^, partly denaturing them^65^. Of the nine PLD-inducers, three (amiodarone, itraconazole, and ivermectin) scattered light intensely by dynamic light scattering (DLS), as did the positive control aggregator, sorafenib^66, 67^ (**Figure S3**). All four molecules also underwent a phase transition to the colloidal form, with critical aggregation concentrations (CAC) in the low µM range (**Figure 5C**) which we note is below their inhibitory concentrations in LPLA2 assay. This is consistent with a colloidal (non-specific) mechanism for LPLA2 inhibition. To further test for this mechanism, we spun down solutions of the four colloid-forming molecules in LPLA2 assay buffer on a benchtop microfuge for 30 minutes. This will pellet-out the ∼200 nM radius colloidal drug particles but leave well-behaved soluble inhibitors in solution^15, 62, 68^. After the spin, we collected the supernatant, added the LPLA2 enzyme and its substrate, and measured activity (**Figure 5D**). If the drug colloids are the active inhibitory species rather than the soluble monomer, the supernatant should have much reduced inhibition. For instance, the well-characterized colloidal aggregator sorafenib, which is not a CAD or a PLD-inducer, potently inhibited LPLA2 before but not after spin-down (**Figure 5D**). Amiodarone and itraconazole saw similar behavior, but ivermectin saw no detectable change in inhibition before and after spin-down. Taken together, these observations demonstrate that seven out of the nine PLD-inducers screened do not inhibiting LPLA2 at concentrations where they robustly induce phospholipidosis. The two that do inhibit the enzyme in a relevant concentration range, amiodarone and itraconazole appear to act via a non-specific mechanism, colloidal aggregation, and are unlikely to be relevant in vivo. These results suggests that inhibition of LPLA2 may be disconnected from PLD-induction and perhaps should not be used as a proxy-assay for it.

## Discussion

Three observations from this study merit particular emphasis. **First**, PLD inducers are prevalent in antiviral drug repurposing screens. Out of 40 putatively antiviral drugs that emerged from these screens and were tested here, 26 (65%) induced phospholipidosis in the low- to sub-micromolar range, overlapping with their reported antiviral EC_50_ activities. This supports the idea that drug-induced phospholipidosis is a general artifact in antiviral drug repurposing, affecting multiple viruses. **Second**, and intriguingly, four of the phospholipidosis inducers were not classical cationic amphiphilic drugs (CADs). Two of the drugs, daptomycin and azithromycin are cationic but are not amphiphilic (**Figure 2C**) and lack the hydrophobic moieties characteristic of CADs. Another two, itraconazole and ivermectin, are not cationic even at lysosomal pH values. Meanwhile, there are both cationic and cationic amphiphilic drugs that do not induce phospholipidosis, such as melperone and elacridar^17^. While most known phospholipidosis inducers are cationic amphiphiles, being a cationic amphiphile seems neither necessary nor sufficient to induce phopholipidosis^51^. Although surprising, this observation aligns with emerging reports from other groups identifying non-CAD compounds as phospholipidosis inducers^61^. These non-ionizable compounds suggest that proposed mechanisms for drug-induced phospholipidosis that depend solely on the titration of ionizable groups into cations^21, 46, 47^ are incomplete. **Third**, our results indicate that the mechanism responsible for PLD induction and its associated antiviral activity differs from previously proposed models involving lysosomal alkalinization or LPLA2 inhibition. A wide range of PLD inducers had little or often no measurable effect on lysosomal pH, as measured by the Fire-pHLy biosensor, suggesting that strong lysosomal alkalinization may not be necessary for phospholipid accumulation nor for antiviral effect. Nor did these PLD inducers inhibit the lysosomal enzyme LPLA2, except via non-specific colloidal aggregation.

These results highlight a broader challenge in drug discovery: lipophilic, bioactive compounds often act through off-target or artifactual mechanisms such as phospholipidosis or colloidal aggregation. While drug induced phospholipidosis cannot be fully predicted based on drug physical properties or known mechanism of action, we describe a facile assay, optimized based on an earlier study^48^, that allows drug-induced phospholipidosis to be rapidly characterized in dose response, ruling out this artifact and allowing investigators to focus on more promising candidates. This assay and strategy may thus join others that have been developed since the advent of high-throughput screening^69–71^ and drug repurposing^15, 72^ to rapidly rule out artifacts in early discovery.

Certain caveats should be mentioned. We only investigated the correlation between phospholipidosis and antiviral activity. We do not claim to fully understand how phospholipidosis leads to an antiviral effect; this merits further study. Additionally, the non-CAD PLD inducers identified here were selected somewhat arbitrarily based on frequent appearances in the antiviral literature and the absence of a well-defined mechanism of action. Meanwhile, drugs like nelfinavir are validated antivirals, even though they are also phospholipidosis inducers. For nelfanivir, the antiviral potency is at least ten-fold lower than its EC_50_ for phospholipidosis induction. We also acknowledge that PLD induction can be cell line dependent. Thus, while recent studies suggest that ivermectin does not induce PLD against SARS-CoV-2 in A549 cells^73^, in HepG2 cells it does so, and that activity may play out as antiviral against other viruses depending on the cell line in which the viral assay is performed. We encourage investigators to screen for phospholipidosis in the cell line most relevant to their system.

Returning to our main theme, a pessimistic outcome of this study is that many drugs repurposed against multiple viruses have cell-based activities that overlap with phospholipidosis induction, and likely have their antiviral actions via this mechanism. Such molecules are unlikely to progress therapeutically^17^ as antivirals. More optimistically, phospholipidosis can rapidly and quantitatively detected by the LipidTox assay used here; counter screening with this assay in early antiviral discovery may save much time and resources. The potent induction of phospholipidosis by drugs like itraconazole and ivermectin, which are distinctly non-CADs, points to a more widespread prevalence of this effect among bioactive molecules. Finally, while we have focused on antiviral drug discovery, the general disruption of cellular lipid homeostasis by phospholipidosis may make it an artifact in drug screens for other diseases as well, not least those affecting the lysosome and mechanisms associated with it.

## Experimental Section

### Literature search

To identify drug candidates to screen for phospholipidosis, we searched Pubmed and Google Scholar with the keywords: “FDA-approved drug”, “antiviral”, “drug repositioning”, and “drug repurposing”. To filter out Covid-19 results, we added common viral names (e.g., ebola, zika, and influenza) for which investigators are still pursuing therapeutics. Identified compounds were active in the mid-nanomolar to micromolar range in cell-based assays. Drug SMILES were generated using ChemDraw version 22.2.0.3348. Using these SMILES, a drug’s cLogP (protonated SMILES pH 7.4) and most basic pKa was calculated using the Chem.Descriptors module from RDkit-2018.09 and JChem-21.13, respectively. Drugs that met the following criteria: cLogP≥ 2, calculated pKa > 7.4, were added to the testing list. If a drug was an already reported phospholipidosis inducer, it was also added to the drug candidates list.

### Compounds

All compounds were supplied as >95% pure by HPLC analysis as reported by the vendors and were used as supplied without further purification. Compounds were ordered from Sigma Aldrich, SelleckChem, Cayman Chemical, TargetMol, or Medchem Express.

### Cell lines

Hep G2 cells (ATCC, HB-8065) were maintained in Eagle’s Minimum Essential Medium (EMEM, Corning, 10-009-CV) supplemented with 10% Fetal Bovine Serum (FBS, Caisson Labs, FBL01) and 1X Penicillin-Streptomycin (Sigma-Aldrich, P4333) and were grown at 37°C and 5% CO_2_. All cells and reagents were guaranteed mycoplasma free by supplier with no additional testing carried out.

### Phospholipidosis assay (LipidTox Red)

Hep G2 cells were seeded at a density of 1 x 10^4^ in black 96-well plate (Corning, 3340) in EMEM media supplemented with 10% FBS and 1X Penicillin-Streptomycin. The following day, media was removed and replaced with fresh media containing test compound with final DMSO concentration 1% (v/v) and 1X LipidTox red dye (Fisher Scientific, H34351). After 24 hours, fluorescence emission was measured at 618nm using CLARIOstar plate reader (BMG Labtech). Following fluorescence reading, nuclei stain (Thermo Fisher Scientific, 62249) was added to cells, incubated for 15 minutes, and fluorescence was measured at excitation/emission of 360/480. Phospholipidosis fluorescence was normalized to cell count by dividing phospholipidosis values by nuclei stain values. All compounds screened were normalized to amiodarone, a well-characterized phospholipidosis inducer. Each compound was tested in triplicate for a total of three independent experiments. Data was analyzed using GraphPad Prism software version 10.1.1 (San Diego, CA).

### Phospholipidosis assay (NBD-PE)

Similar to LipidTox Red assay, Hep G2 cells were seeded at a density of 1 x 10^4^ in black 96-well plate in EMEM media supplemented with 10% FBS and 1X Penicillin-Streptomycin. The following day, media was removed and replaced with fresh media containing test compound with final DMSO concentration 0.2% (v/v) and 7.5µM NBD-PE (ThermoFisher, N360). Cells were incubated for another 24 hours before nuclei staining and fixation with 4% (v/v) paraformaldehyde. Images were taken on a CellInsight CX7 (ThermoFisher) equipped with a 20x objective. Drug conditions were performed in triplicate with 9-fields taken from each well. Images were analyzed using the HCS Studio software (ThermoFisher).

### Dynamic Light Scattering (DLS)

Compounds were diluted in filtered 50 mM KPi buffer, pH 7, at a final concentration of 1% DMSO (v/v). All compounds were initially screened at a top concentration of 100 µM using a Wyatt DynaPro Plate Reader II (Waters Corporation). Samples that had a scattering intensity > 1×107 cnts/s or 1-fold over baseline scattering were considered to be forming colloidal-like particles (50-1000 nm diameter) and were re-screened as a concentration-response in eight-point half-log dilutions. During data analysis, data were separated into two groups: aggregating concentrations (scattering > 1×10^7^ cnts/s) and non-aggregating concentrations (scattering < 1×10^7^ cnts/s). A line was generated for each group and the point of intersection of the two lines serves as the critical aggregation concentration (CAC).

### LPLA2 Enzyme Inhibition Assay

Compounds were screened using a commercial kit (Echelon Biosciences, K-70001) following manufacturer instruction. Reagent grade water was added to LPLA2 substrate followed by vortexing for 5 minutes to form liposomes. LPLA2 substrate and test compounds were pre-incubated for 1 hour RT. After pre-incubation, LPLA2 enzyme and reaction buffer (1X final) were added and incubated an additional hour at RT with shaking. All compounds were screened at a final concentration of 31.6 uM with final DMSO concentration 6.25% (v/v). A DMSO only, no-compound control was added to measure baseline enzyme activity, and a no-enzyme sample was used to measure background. After a 1-hour incubation, the reaction was stopped by adding stop buffer (1X final) and measured using a CLARIOStar Plus Plate Reader (BMG Biotech). Fluorescence was recorded at 490 nm excitation/540 nm emission and background subtracted from raw values. All data were then normalized with the DMSO-only control sample indicating 100% enzyme activity.

To rule out the effects of aggregation, a centrifugation step was added. In this modified procedure, the compound is first added to reaction buffer (1X final) and centrifuged using a Sorvall Legend Micro 21R centrifuge (Thermo Scientific) at 15,700 xg for 30 minutes to remove colloidal particles^63^. For method validation, this modified procedure with and without the centrifugation steps were compared to samples prepared using the original kit instructions. The data for each procedure were normalized to their respective DMSO-only control sample. After method validation, all compounds displaying greater than 30% LPLA2 inhibition using the original protocol were re-screened with the centrifugation procedure to mitigate the effects of aggregation.

## ASSOCIATED CONTENT

### Supporting information

Supporting Information is available free of charge on the ACS Publication website.

- Table showing all compounds tested, concentration-response for additional repurposed drug candidates that induce phospholipidosis, and single-point concentration of repurposed drug candidates that did not induce phospholipidosis

## Author Contributions

Conceived by BKS, ISG and ADW. Proof of concept assays done by ADW; all results reported here by ISG. LP screening compounds for phospholipidosis. MM, MMK, and MO intellectual contributions to virology.

## Notes

The authors declare the following competing financial interest(s): BKS is co-founder of BlueDolphin LLC, Epiodyne Inc, and Deep Apple Therapeutics, Inc., and serves on the SRB of Genentech, the SAB of Schrodinger LLC, and the SAB of Vilya Therapeutics. No other authors declare competing interests.

## Abbreviation Used

CAD: Cationic amphiphilic drug
DIPL: Drug-induced Phospholipidosis
NBD-PE: N-(7-Nitrobenz-2-Oxa-1,3-Diazol-4-yl)-1,2-Dihexadecanoyl-sn-Glycero-3-Phosphoethanolamine Triethylammonium Salt
PLD: Phospholipidosis
SD: Standard Deviation

## TOC Graphic

### For Table of Contents Only

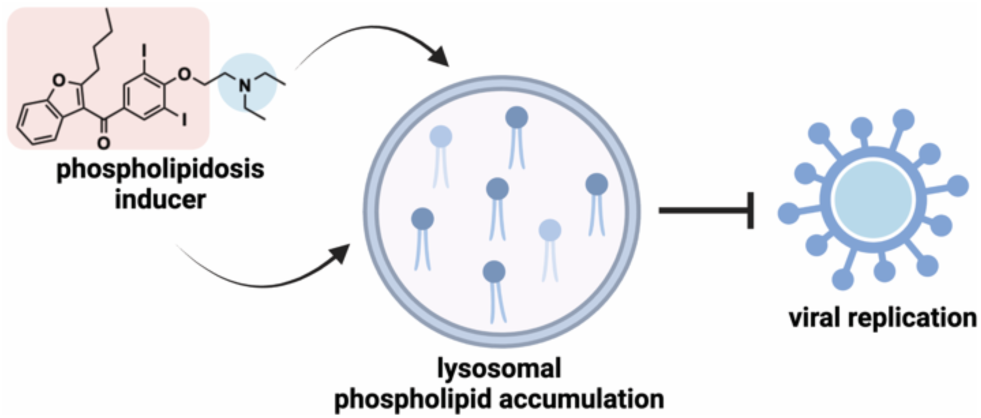

