## Supporting Information for "Drug-induced phospholipidosis as an artifact in antiviral drug repurposing"

**Affiliations:**

**Contents of SI:**

1. Table showing all compounds tested
2. Concentration-response for additional repurposed drug candidates that induce phospholipidosis
3. Single-point concentration of repurposed drug candidates that did not induce phospholipidosis
4. One-way ANOVA statical test of FIREpHLy ratio fold-change
5. Dynamic light scattering of PLD inducers

Table 1. Phospholipidosis candidates

| Name of drug | SMILES | Reported target | Literature reference (PMID) | **cLogP** | **pKa (basic)** | **Vendor** | **Catalog number** |
|---|---|---|---|---|---|---|---|
| Amiodarone | IC1=C(OCCN(CC)CC)C(I)=CC(C(C2=C(CCCC)OC3=C2C=CC=C3)=O)=C1 | EBOV, HCV, MARV | 30016444; 19995961; 24710028 | 5.5191 | 9.08 | Cayman Chemical Company | 15213 |
| Amodiaquine | CCN(CC1=C(C=CC(NC2=C3C=CC(Cl)=CC3=NC=C2)=C1)O)CC | DENV2, ZIKV | 24680954; 29315671 | 3.7621 | 10.28 | Cayman Chemical Company | 15954 |
| Aripiprazole | ClC1=CC=CC(N2CCN(CCCCOC3=CC=C(CCC(N4)=O)C4=C3)CC2)=C1Cl | EBOV | 26041706 | 3.4422 | 7.46 | Sigma-Aldrich | SML0935 |
| Azithromycin | CC[C@H]1OC([C@@H]([C@H]([C@@H]([C@H]([C@](O)(C[C@H](CN([C@@H]([C@H]([C@@]1(O)C)O)C)C)C)C)O[C@@H]2O[C@@H](C[C@H](N(C)C)[C@H]2O)C)C)O[C@H]3C[C@@](OC)([C@H]([C@@H](O3)C)O)C)C)=O | DENV2,IAV, ZIKV | 31527024; 31300721; 31527024 | -0.9335 | 9.68 | Cayman Chemical Company | 15004 |
| Benidipine | COC(C1=C(NC(C)=C(C(O[C@@H]2CCCN(C2)CC3=CC=CC=C3)=O)[C@@H]1C4=CC=CC([N+]([O-])=O)=C4)C)=O | JEV, ZIKV | 28814523 | 2.7933 | 8.09 | Cayman Chemical Company | 21607 |
| Bepridil | CC(C)COCC(CN(CC1=CC=CC=C1)C2=CC=CC=C2)N3CCCC3 | EBOV | 26041706 | 3.4131 | 9.16 | MedChem Express | HY-103315 |
| Berbamine | NC1=C2C3CC4=C(C=CC=C4)CN3CCC2=CC=C1 | JEV | 34102949 | 2.9243 | 6.95 | Selleck Chemicals | S3609 |
| Cilnidipine | COCCOC(C1=C(NC(C)=C(C(OC/C=C/C2=CC=CC=C2)=O)C1C3=CC=CC([N+]([O-])=O)=C3)C)=O | JEV, ZIKV | 28814523 | 4.2758 | -4.12 | MedChem Express | HY-17404 |

| | | | | | | | |
|---|---|---|---|---|---|---|---|
| Daptomycin | O=C(CCCCCCCCC C)N[C@H](C(N[C @@H](CC(N)=O) C(N[C@@H](CC( O)=O)C(NC(C(OC ([C@@H](NC([C @@H](NC(C(NC( CNC([C@@H](N C1=O)CC(O)=O)= O)=O)CO)=O)[C @H](CC(O)=O)C) =O)CC(C2=C(N)C =CC=C2)=O)=O) C)C(NCC(N[C@H ](CCCN)C(NC(CC (O)=O)C(N[C@@ H]1C)=O)=O)=O)= O)=O)=O)=O)CC3 =CNC4=C3C=CC =C4 | ZIKV | 27476412 | -11.677 | 9.59 | Selleck Chemicals | S1373 |
| Doxepin | CN(C)CC/C=C1C 2=C(C=CC=C2)O CC3=C/1C=CC=C 3 | HCV | 19995961 | 2.5453 | 9.76 | Cayman Chemical Company | 15888 |
| Dronedarone | CCCCN(CCCOC1 =CC=C(C(C2=C( OC3=C2C=C(C= C3)NS(C)(=O)=O) CCCC)=O)C=C1) CCCC | EBOV, MARV | 24710028 | 5.6319 | 10.31 | MedChem Express | HY-A0016 |
| Emetine | CC[C@@H]1[C@ @H](C[C@H]2NC CC3=C2C=C(OC) C(OC)=C3)C[C@ H]4C5=C(C=C(C( OC)=C5)OC)CCN 4C1 | DENV2 | 24941340 | 2.5001 | 9.23 | Cayman Chemical Company | 21048 |
| Itraconazole | CCC(C)N1C(N(C= N1)C2=CC=C(N3 CCN(CC3)C4=CC =C(OC[C@H]5CO [C@](C6=C(Cl)C= C(Cl)C=C6)(O5)C N7C=NC=N7)C=C 4)C=C2)=O | EBOV | 35337896 | 5.5773 | 3.91 | Selleck Chemicals | S2476 |

| | | | | | | | |
|---|---|---|---|---|---|---|---|
| Ivermectin | C/C1=C\C[C@@H](O2)C[C@@H](C[C@@]32CC[C@H](C)[C@]([C@@H](C)CC)([H])O3)OC([C@@H]([C@]([C@]45[H])(O)/C(CO4)=C/C=C/[C@@H]([C@@H]1O[C@@H]6O[C@@H](C)[C@H](O[C@@H]7O[C@@H](C)[C@H](O)[C@@H](OC)C7)[C@@H](OC)C6)C)C=C(C)[C@H]5O)=O | ZIKV | 27476412 | 5.6014 | -3.45 | Sigma-Aldrich | I8898 |
| Manidipine | COC(C1=C(NC(C)=C(C(OCCN2CCN(C(C3=CC=CC=C3)C4=CC=CC=C4)CC2)=O)C1C5=CC=CC([N+]([O-])=O)=C5)C)=O | JEV, ZIKV | 28814523 | 3.5359 | 7.89 | MedChem Express | HY-17403 |
| Memantine | CC12CC3CC(C1)(CC(C3)(C2)N)C | ZIKV, CoV | 29759926; 24227863 | 1.9773 | 10.7 | Sigma-Aldrich | M9292 |
| Nelfinavir | [H][C@@]12CCCC[C@@]1(CN([C@H](C(NC(C)(C)C)=O)C2)C[C@H]([C@@H](NC(C3=C(C(O)=CC=C3)C)=O)CSC4=CC=CC=C4)O)[H] | HCV, JEV, ZIKV | 19995961; 28814523 | 3.33052 | 8.18 | Cayman Chemical Company | 15144 |
| Orphenadrine | CN(CCOC(C1=CC=CC=C1C)C2=CC=CC=C2)C | EBOV, MARV | 29981374 | 2.24552 | 8.87 | Cayman Chemical Company | 26391 |
| Quinacrine | CCN(CCCC(NC1=C2C=C(C=CC2=NC3=C1C=CC(Cl)=C3)OC)C)CC | EBOV | 26041706 | 3.9744 | 10.33 | Selleck Chemicals | S5435 |
| Sertraline | CN[C@H]1CC[C@H](C2=CC=CC=C21)C3=CC(Cl)=C(Cl)C=C3 | EBOV | 26041706 | 4.1534 | 9.56 | Cayman Chemical Company | 14839 |
| Terfenadine | CC(C)(C1=CC=C(C(CCCN2CCC(C(C3=CC=CC=C3)(O)C4=CC=CC=C4)CC2)O)C=C1)C | HCV | 18560330 | 5.0287 | 9.02 | Selleck Chemicals | S4353 |
| Thioridazine | CSC1=CC2=C(C=C1)SC3=CC=CC=C3N2CCC4CCCCN4C | EBOV | 26041706 | 4.4685 | 8.93 | MedChem Express | HY-B0965 |
| Tilorone | CCN(CCOC1=CC=C2C(C(C3=C2C=CC(OCCN(CC)CC)=C3)=O)=C1)CC | EBOV | 26834994 | 1.505 | 9.49 | Cayman Chemical Company | 17868 |

| | | | | | | | |
|---|---|---|---|---|---|---|---|
| Toremifene | ClCC/C(C1=CC=CC=C1)=C(C2=CC=CC=C2)/C3=CC=C(OCCN(C)C)C=C3 | EBOV, HCV | 26041706; 19995961 | 4.7979 | 8.76 | Cayman Chemical Company | 20854 |
| U18666A | C[C@]1([C@](CC2)([H])[C@]3([H])CC=C4C[C@@H](OCCN(CC)CC)CC[C@]4(C)[C@@]3([H])CC1)C2=O.Cl | EBOV, DENV | 22146564; 23441171 | 3.8282 | 9.41 | MedChem Express | HY-107433 |
| Verapamil | N#CC(CCCN(C)CCC1=CC(OC)=C(OC)C=C1)(C2=CC(OC)=C(OC)C=C2)C(C)C | EBOV, IAV, MARV | 24710028; 6743023 | 3.67598 | 9.68 | Cayman Chemical Company | 14288 |
| Acyclovir | OCCOCN(C=N1)C2=C1C(NC(N)=N2)=O | HSV, CoV-2 | 37228547 | -1.3318 | 0.6 | Sigma-Aldrich | 1012065 |
| Carbinoxamine maleate | CN(CCOC(C1=CC=CC=N1)C2=CC=C(C=C2)Cl)C.O=C(/C=C\C(O)=O)O | IAV | 30459739 | 1.9855 | 8.87 | TargetMol | T2222 |
| Chlorpheniramine maleate | CN(C)CCC(C1=NC=CC=C1)C2=CC=C(Cl)C=C2.O=C(O)/C=C\C(O)=O | IAV | 30459739 | 2.4015 | 9.47 | Cayman Chemical Company | 21253 |
| Favipiravir | O=C(C1=NC(F)=CN=C1O)N | EBOV | 24462697; 11897578 | -0.5798 | -3.68 | TargetMol | T6833 |
| Indinavir | CC(C)(NC([C@@H]1CN(CCN1C[C@H](C[C@H](C(N[C@@H]2[C@@H](CC3=CC=CC=C23)O)=O)CC4=CC=CC=C4)O)CC5=CN=CC=C5)=O)C | EBOV | 26887654 | 2.8669 | 6.76 | Cayman Chemical Company | 15150 |
| Mycophenolic acid | COC1=C(C(O)=C2C(OCC2=C1C)=O)C/C=C(CCC(O)=O)\C | ZIKV | 29315671 | 1.39852 | -4.07 | Sigma-Aldrich | M5255 |
| Nifedipine | O=C(C1=C(C)NC(C)=C(C(OC)=O)C1C2=CC=CC=C2[N+]([O-])=O)OC | SFTSV | 31444469 | 2.1756 | -6.64 | Sigma-Aldrich | N7634 |
| Nitazoxanide | O=C(NC1=NC=C([N+]([O-])=O)S1)C2=CC=CC=C2OC(C)=O | IAV | 19995961 | 2.2289 | -4.25 | Cayman Chemical Company | 13692 |
| Oseltamivir | CCOC(C1=C[C@H]([C@@H]([C@H](C1)N)NC(C)=O)OC(CC)CC)=O | IAV, CoV-2 | 36454880 | 0.5686 | 9.26 | Sigma-Aldrich | SML1606 |
| Pioglitazone | CCC1=CN=C(C=C1)CCOC2=CC=C(C=C2)CC3SC(NC3=O)=O | HCV | 22412837 | 3.9437 | 5.63 | Cayman Chemical Company | 10028 |

| | | | | | | | |
|---|---|---|---|---|---|---|---|
| Ribavirin | OC[C@@H]1[C@H]([C@H]([C@H](N2C=NC(C(N)=O)=N2)O1)O)O | HCV | 21960835 | -3.0115 | -1.16 | Cayman Chemical Company | 16757 |
| Ruxolitinib | N#CC[C@@H](N1C=C(C2=C3C=CNC3=NC=N2)C=N1)C4CCCC4 | HIV | 24419350 | 3.46638 | 3.91 | Selleck Chemicals | S1378 |
| Sofosbuvir | CC(OC([C@@H](N[P@](OCC1O[C@H]([C@](F)([C@@H]1O)C)N2C=CC(NC2=O)=O)(OC3=CC=CC=C3)=O)C)=O)C | ZIKV | 24983590 | 1.6565 | -3.85 | MedChem Express | HY-15005 |
| Zanamivir | N[C@H]1C=C(C(OC)=O)O[C@]([C@@H]([C@@H](COC(C)=O)OC(C)=O)OC(C)=O)([H])[C@@H]1NC(C)=O | IAV | 9302301 | -2.0162 | 8.66 | Cayman Chemical Company | 15123 |

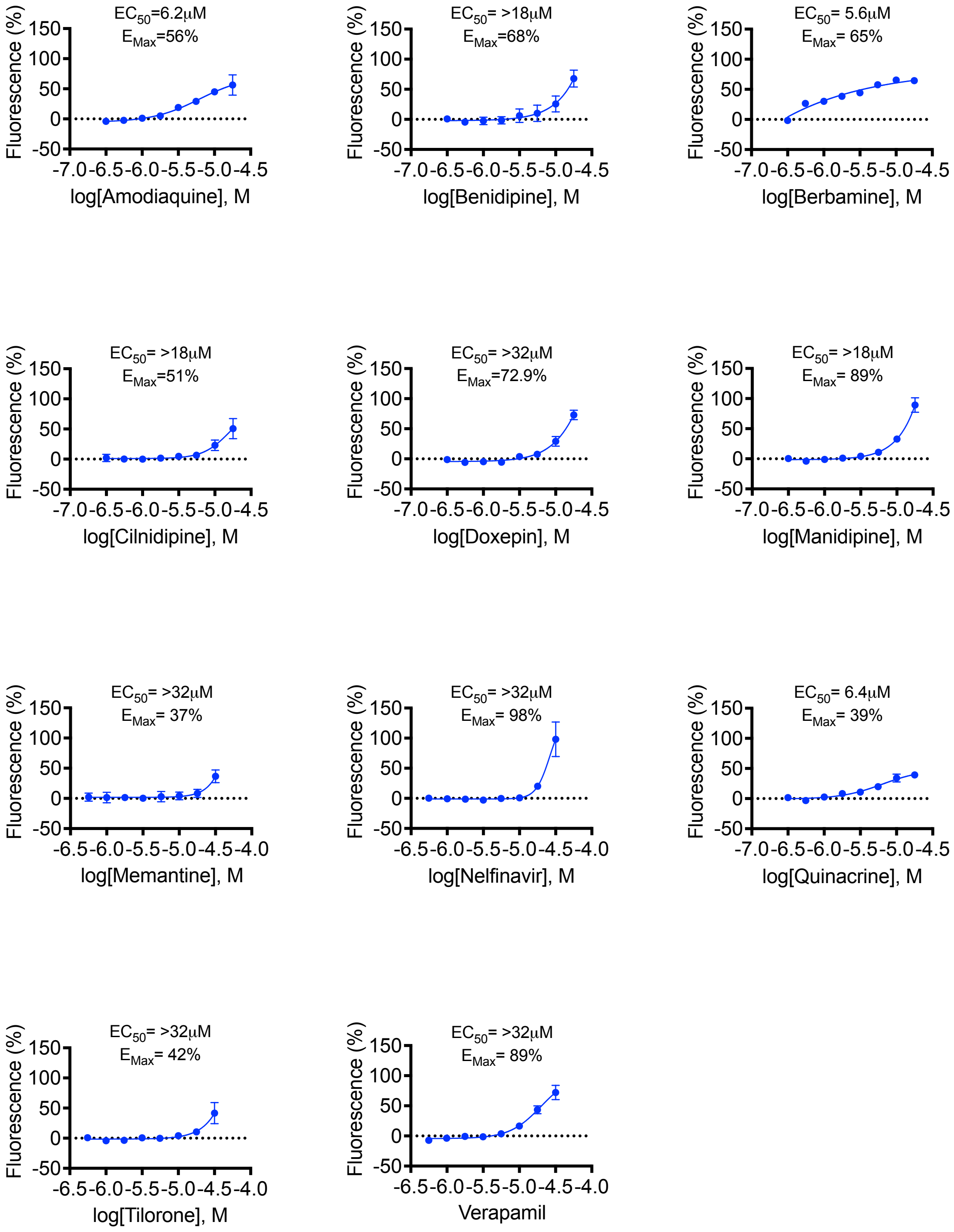


**Figure S1.** Additional repurposed drug candidates that induce phospholipidosis. Data shown was performed in HepG2 cells using LipidTox Red phospholipid stain for three independent experiments performed in triplicate. Data was normalized to positive control compound, amiodarone, and cell count using Hoechst nuclei stain.

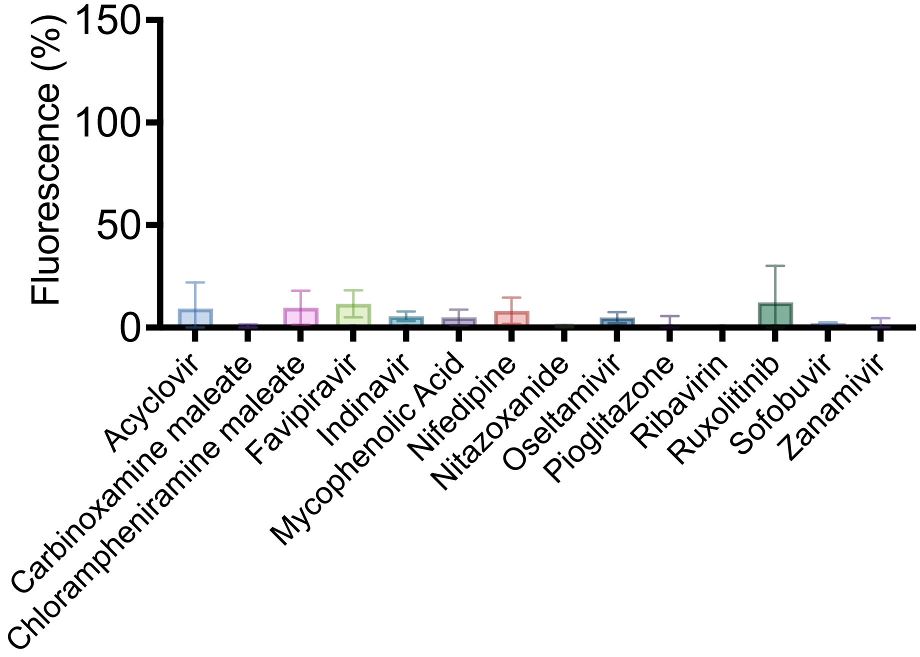


**Figure S2.** Repurposed drug candidates that did not induce phospholipidosis. Single-point concentration treatment of drug (31.6uM). All data shown was performed in HepG2 cells using LipidTox Red phospholipid stain for a single independent experiment performed in triplicate. Data was normalized to positive control compound, amiodarone, and cell count using Hoechst nuclei stain.

| Bonferroni's multiple comparisons test | Mean Diff. | 95.00% CI of diff. | Below threshold? | Summary | Adjusted P Value |
|---|---|---|---|---|---|
| DMSO - 0.1% vs. DMSO - 0.18% | 0.00 | -0.1355 to 0.1355 | No | ns | >0.9999 |
| DMSO - 0.1% vs. Water - 0.1% | 0.00 | -0.1355 to 0.1355 | No | ns | >0.9999 |
| DMSO - 0.1% vs. Water - 0.18% | 0.00 | -0.1355 to 0.1355 | No | ns | >0.9999 |
| DMSO - 0.1% vs. BaF (100 nM) | -2.46 | -2.575 to -2.340 | Yes | **** | <0.0001 |
| DMSO - 0.1% vs. BaF (100 nM) | -2.46 | -2.575 to -2.340 | Yes | **** | <0.0001 |
| DMSO - 0.18% vs. Amiodarone (17.8 µM) | -0.14 | -0.2905 to 0.002282 | No | ns | 0.0581 |
| DMSO - 0.1% vs. Azithromycin (10 µM) | -0.39 | -0.5314 to -0.2386 | Yes | **** | <0.0001 |
| DMSO - 0.18% vs. Azithromycin (17.8 µM) | -0.57 | -0.7127 to -0.4199 | Yes | **** | <0.0001 |
| DMSO - 0.1% vs. Carbinoxamine maleate (10 µM) | -0.01 | -0.1517 to 0.1411 | No | ns | >0.9999 |
| DMSO - 0.18% vs. Carb. maleate (17.8 µM) | -0.01 | -0.1539 to 0.1389 | No | ns | >0.9999 |
| DMSO - 0.1% vs. Doxepin (10 µM) | 0.04 | -0.1018 to 0.1910 | No | ns | >0.9999 |
| DMSO - 0.18% vs. Doxepin (17.8 µM) | 0.05 | -0.09474 to 0.1980 | No | ns | >0.9999 |
| Water - 0.1% vs. Emetine (10 µM) | -0.23 | -0.3723 to -0.07956 | Yes | *** | 0.0001 |
| Water - 0.18% vs. Emetine (17.8 µM) | -0.33 | -0.4772 to -0.1844 | Yes | **** | <0.0001 |
| DMSO - 0.1% vs. Itraconazole (1.78 µM) | -0.02 | -0.1638 to 0.1290 | No | ns | >0.9999 |
| DMSO - 0.1% vs. Itraconazole (3.16 µM) | -0.11 | -0.2580 to 0.03482 | No | ns | 0.4246 |
| DMSO - 0.1% vs. Ivermectin (10 µM) | -0.07 | -0.2153 to 0.07745 | No | ns | >0.9999 |
| DMSO - 0.18% vs. Ivermectin (17.8 µM) | -0.44 | -0.5903 to -0.2975 | Yes | **** | <0.0001 |
| DMSO - 0.1% vs. Melperone (10 µM) | -0.06 | -0.2014 to 0.09139 | No | ns | >0.9999 |
| DMSO - 0.18% vs. Melperone (17.8 µM) | -0.10 | -0.2433 to 0.04951 | No | ns | 0.9362 |
| DMSO - 0.1% vs. Terfenadine (10 µM) | -0.19 | -0.3360 to -0.04318 | Yes | ** | 0.0023 |
| DMSO - 0.18% vs. Terfenadine (17.8 µM) | -0.58 | -0.7256 to -0.4328 | Yes | **** | <0.0001 |
| DMSO - 0.1% vs. Toremifene (10 µM) | 0.01 | -0.1380 to 0.1548 | No | ns | >0.9999 |
| DMSO - 0.18% vs. Toremifene (17.8 µM) | -0.15 | -0.3010 to -0.008217 | Yes | * | 0.0287 |
| DMSO - 0.1% vs. U18666A (1.78 µM) | 0.02 | -0.1269 to 0.1659 | No | ns | >0.9999 |
| DMSO - 0.1% vs. U18666A (3.16 µM) | -0.03 | -0.1740 to 0.1188 | No | ns | >0.9999 |

**Table S2.** One-way ANOVA statical test of FIREpHLy ratio fold-change in HepG2 cells after 24 hr drug treatment. Data points are from three technical replicates.

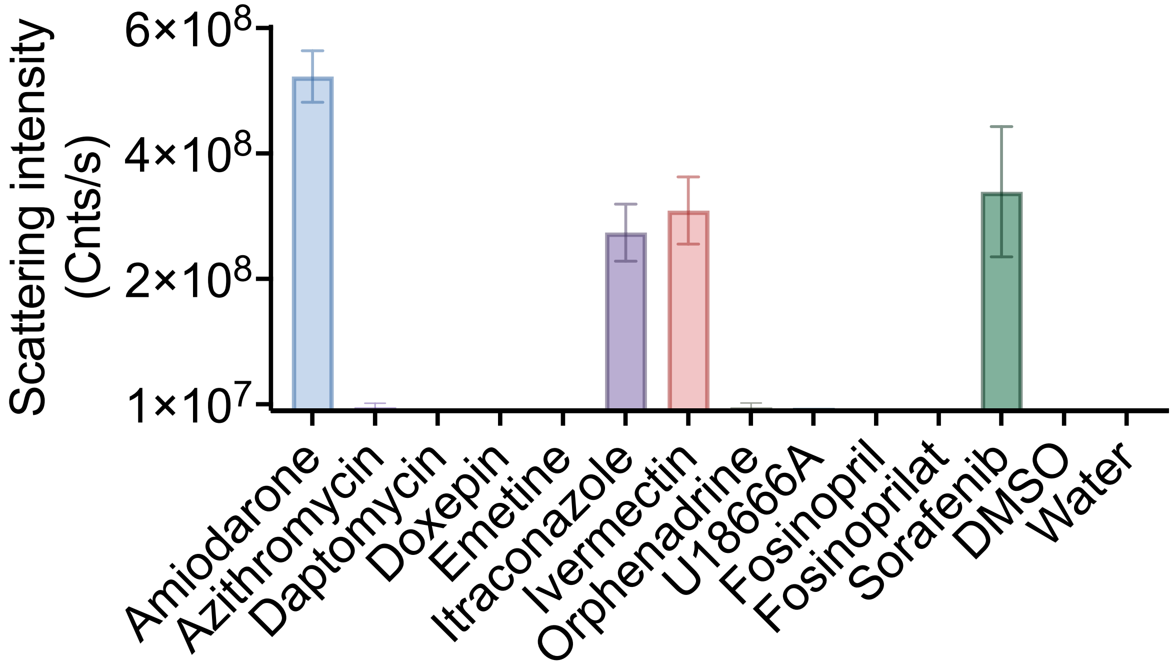


**Figure S3.** Dynamic light scattering performed on colloidal aggregator candidates. Compounds with scattering intensity greater than $1x10^7$ or 1-fold over baseline scattering and particles size of 50-1000 nm in diameter were considered colloidal aggregators. All compounds were screened at 100 μM. Data points are presented as mean ± SD from three technical replicates.
